# Dilated Convolutions for Modeling Long-Distance Genomic Dependencies

**DOI:** 10.1101/200857

**Authors:** Ankit Gupta, Alexander M. Rush

**Affiliations:** School of Engineering and Applied Sciences, Harvard University, Cambridge, MA 02138

## Abstract

We consider the task of detecting regulatory elements in the human genome directly from raw DNA. Past work has focused on small snippets of DNA, making it difficult to model long-distance dependencies that arise from DNA’s 3-dimensional conformation. In order to study long-distance dependencies, we develop and release a novel dataset for a larger-context modeling task. Using this new data set we model long-distance interactions using dilated convolutional neural networks, and compare them to standard convolutions and recurrent neural networks. We show that dilated convolutions are effective at modeling the locations of regulatory markers in the human genome, such as transcription factor binding sites, histone modifications, and DNAse hypersensitivity sites.

## 1 Introduction

Gene expression is controlled by a variety of *regulatory factors* that determine which genes are expressed in which environmental conditions (Perkins et al., 2005). Due to proteins called histones that DNA winds around, parts of DNA are more accessible to binding than others, and so DNA accessibility is a regulatory factor. Modifications to histones can affect the conformation of DNA, so histone modifications are also regulatory factors. Furthermore, proteins that bind to DNA and affect transcriptional activity are called transcription factors and are also regulatory factors. The activity that they affect can be thousands of base pairs away (Blackwood and Kadonaga, 1998).

These interactions imply that nucleotides far apart in a 1-dimensional DNA sequence may interact in its 3-dimensional conformation, and so expression is governed by both local and long-distance dependencies. As a result, it may be important to incorporate DNA regions that are far away in 1-D space when modeling regulatory markers. However, past data sets for binding site prediction have used small snippets of DNA (Alipanahi et al., 2015; Zhou and Troyanskaya, 2015; Quang and Xie, 2016), which limits the ability to model these interactions.

This work addresses this question by introducing a new dataset for this problem allowing for long-distance interactions, and a new model using dilated convolutions to predict the locations of regulatory markers. We learn a mapping from a DNA region, specified as a sequence of nucleotides, to the locations of regulatory markers in that region. Dilated convolutions can capture a hierarchical representation of a much larger input space than standard convolutions, allowing them to scale to large context sizes. We compare dilated convolutions to other modeling methods from deep learning and past work, including standard convolutions and recurrent neural networks, and show that they present an advancement over existing models. All code, data, and scripts for these experiments are available at https://github.com/harvardnlp/regulatory-prediction.

## 2 Models

### Convolutional and Recurrent Neural Networks

The two most widely applied models for sequence tasks are convolutional neural networks (CNNs) and recurrent neural networks (RNNs). CNNs consist of a series of layers each applying a linear transformation followed by a non-linearity to a sliding kernel window of inputs (LeCun et al., 1995), shown in Figure 1(a). RNNs apply the same to each element of a sequence in time. An RNN with state size *M* processes a sequence of inputs {*x*_*i*_}_*N*_ by updating a state vector *c*_*i*_ ∈ ℝ^*M*^ for each timestep *i* such that *c*_*i*_+_1_ = *R*(*c*_*i*_, *x*_*i*_). Concatenating a forward and reverse RNN gives a bidirectional RNN, as in Figure 1(b).

**Figure 1:**
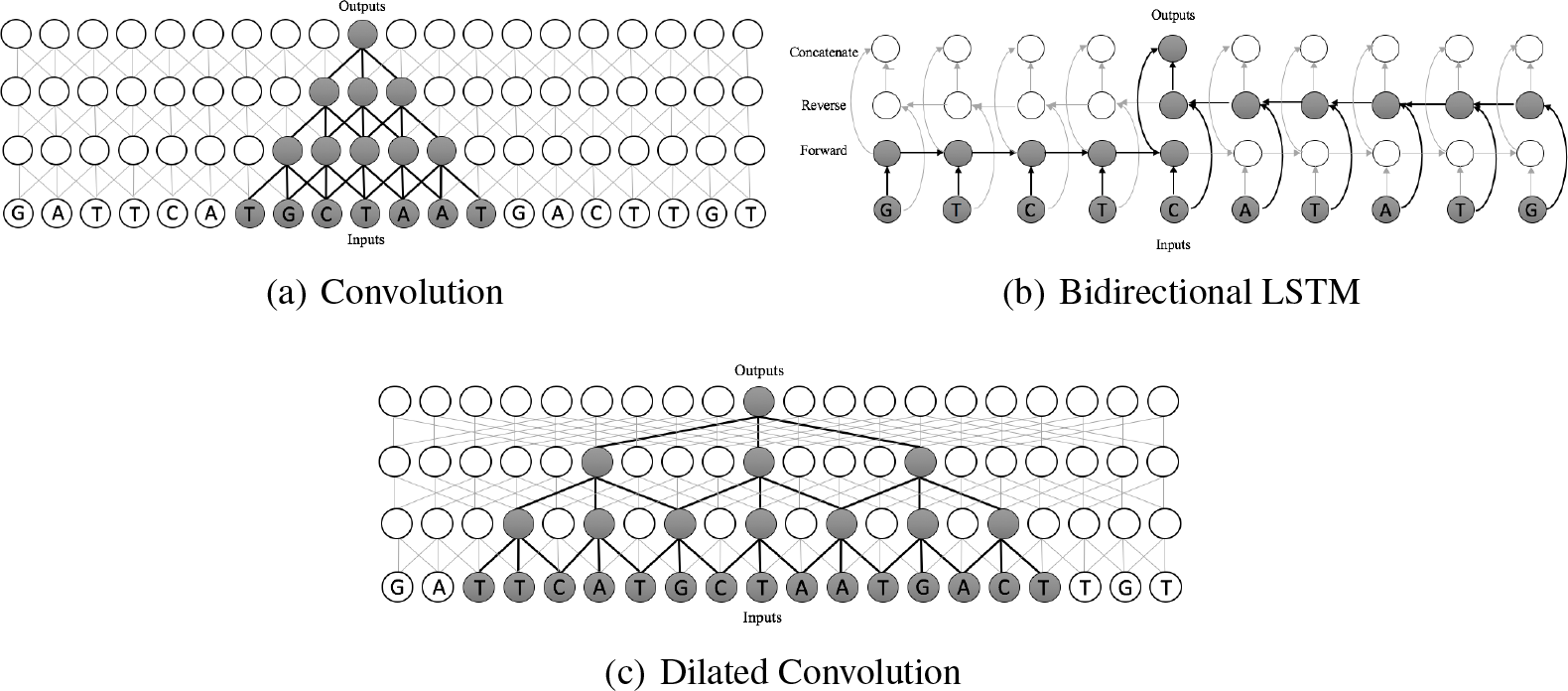
Visual representation of the models. All take a sequence of nucleotides as input. The receptive field is shaded. The convolution has a short path between inputs and predictions but a small receptive field. The Bi-LSTM has a large receptive field, the whole sequence, but may have a long path between a nucleotide and prediction. The dilated convolution has both a short path from the input *and* a large receptive field.

The interactions in a network can be quantified by the receptive field. Formally the *receptive field* of a node is the subset 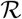 of input elements {*x*_*i*_} that can impact its value (Yu and Koltun, 2015). For CNNs, the size of the receptive field is linear in the number of layers and the kernel width. Thus, scaling the receptive field to incorporate a large input introduces more layers, making training more difficult. Bidirectional RNNs on the other hand have a receptive field of the whole input, but require gradients to travel long-distances over time. Long Short-Term Memory (LSTM) cells (Hochreiter and Schmidhuber, 1997) circumvent some of these issues by using trainable gated connections that allow the cell state to propagate more efficiently. However, with very long sequences, as is the case in genetic data, LSTMs still have trouble learning very long-distance relationships.

### Dilated Convolutions

Dilated convolutions (Yu and Koltun, 2015) offer a middle ground between these two models with wide receptive fields and short distance gradient propagation. In CNNs each kernel window consists of adjacent inputs, while dilated convolutions introduce gaps (dilation) between inputs. With dilation *d*, the window starting at location *i* of size *k* is

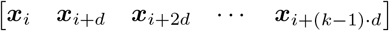

Yu and Koltun (2015) show that by stacking these convolutions with increasingly large *d*, we can expand the receptive field of each output exponentially. This allows them to have large receptive fields, but still short backpropagations, as shown in Figure 1(c). We take advantage of this structure when modeling genetic regulation. Dilated convolutions have been used for image segmentation (Yu and Koltun, 2015), text-to-speech (Oord et al., 2016), and text classification (Strubell et al., 2017).

## 3 Experiments

### Dataset 1: Short Sequence Prediction Benchmark (Zhou and Troyanskaya, 2015)

As a preliminary experiment, we test dilated convolutions on a standard benchmark task and compare them to CNNs and LSTMs. We use the Zhou and Troyanskaya (2015) dataset to predict the presence of regulatory markers in short DNA sequences. Each input is a vector 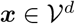, where *d* = 1000 is the sequence length, and 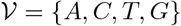. Each output vector *y* ∈ {0,1}^*k*^ indicates whether each of the *k* = 919 regulatory markers is present in the middle 200bp of *x*. The markers include TFBSs, histone modifications, and DNAse hypersensitivity sites (accessible DNA regions).

We train several architectures and report the mean PR AUC scores for each category in Table 1. The models in this task have a final fully-connected layer, which implies that the receptive field of every output contains the whole input (1000 bp). The differences between models is explained by the extent to which each captures meaning prior to the final layer. Note that there was dropout (Srivastava et al., 2014) and batch normalization (Ioffe and Szegedy, 2015) between every layer, and we select hyperparameters using grid search. Details are given in the Supplemental Information section.

**Table 1:** Models and Test Set Precision-Recall Area Under Curve (PR AUC) scores for Dataset 1. We report the number of parameters in the hyperparameter configuration selected using grid search with a held-out validation set. CNN3 is the model from Zhou and Troyanskaya (2015), and Bi-LSTM is the bidirectional LSTM model from Quang and Xie (2016). Our DILATED6 model performs better than the standard convolutions on all three types of predictions and only slightly underperforms the bidirectional LSTM model. All scores are based on reimplementations. Model size varied across hyperparameter configurations.

| Dataset 1: Short Sequence Prediction Task |  |  |  |  |  |  |
| --- | --- | --- | --- | --- | --- | --- |
| Model | Hidden | Type | Params (Best-Case) | Test Set PR AUC |  |  |
|  |  |  |  | TFBS | Hist | DNase |
| LR | 0 | - | 3676919 | 0.042 | 0.143 | 0.097 |
| FF | 1 | Fully | 4551919 | 0.046 | 0.181 | 0.106 |
| CNN3 | 3 | CNN | 155159839 | 0.205 | 0.273 | 0.319 |
| DILATED3 | 3 | Dilated | 37056519 | 0.255 | 0.293 | 0.372 |
| DILATED6 | 6 | Dilated | 25758079 | 0.285 | 0.320 | 0.396 |
| Bi-LSTM | 2 | Bi-LSTM | 46926479 | **0.305** | **0.340** | **0.407** |

Notably the DILATED6 model performs much better than the CNN3 model from Zhou and Troyanskaya (2015), and approaches the performance of the BI-LSTM model. The BI-LSTM is the most effective model, which is reasonable since the sequences are short. This gives a proof-of-concept that dilated convolutions can capture nucleotide structure better than standard convolutions.

### Dataset 2: Complete Genome Labeling with Long-Range Inputs

To model long-distance dependencies, we introduce a new dataset that models the problem as long-distance sequence labeling instead of prediction. In particular: (1) the input sequences are longer, with each a *d* = 25000 bp sequence, and (2) the outputs annotate at nucleotide-resolution, making this a dense labeling task. We predict the presence of all regulatory markers at each nucleotide, rather than per 200bp.

We use hg19 to extract input sequences and *k* = 919 regulatory marker locations from ENCODE (Consortium et al., 2012). Thus, we get pairs of input and output sequences {(*x*, *y*)} with 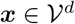and *y* ∈ {0,1}^*d*×*k*^. The inputs have vocabulary 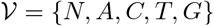, where the N character represents nucleotides with high uncertainty. We remove sequences with >10% unknown nucleotides or with >10% part of a multi-mapped sequence, meaning part of a region that maps to several genomic locations. This left *n* = 93880 non-overlapping sequences that were *d* = 25000 in length, totaling 2.3 billion nucleotides. We train on 80% of the data, and split the rest into validation and test sets.

We train models representing the various architectures on this new dataset. We showed above with Dataset 1 that LSTM-based models were the most effective at predicting regulatory marker locations when given only a small number of nucleotides, and we can now study their relative success when given more context. Since this is a base-pair level prediction task, we no longer have a fully-connected layer for each model. We summarize the models for this task and report PR AUC scores in Table 2. We report the best models across all hyperparameters, including number of filters, dropout, learning rate, batch norm decay, and hidden layer size, using grid search on a held-out validation set. For the dilated convolution, we use dilations of 1, 3, 9, 27, and 81 in sequence.

**Table 2:**
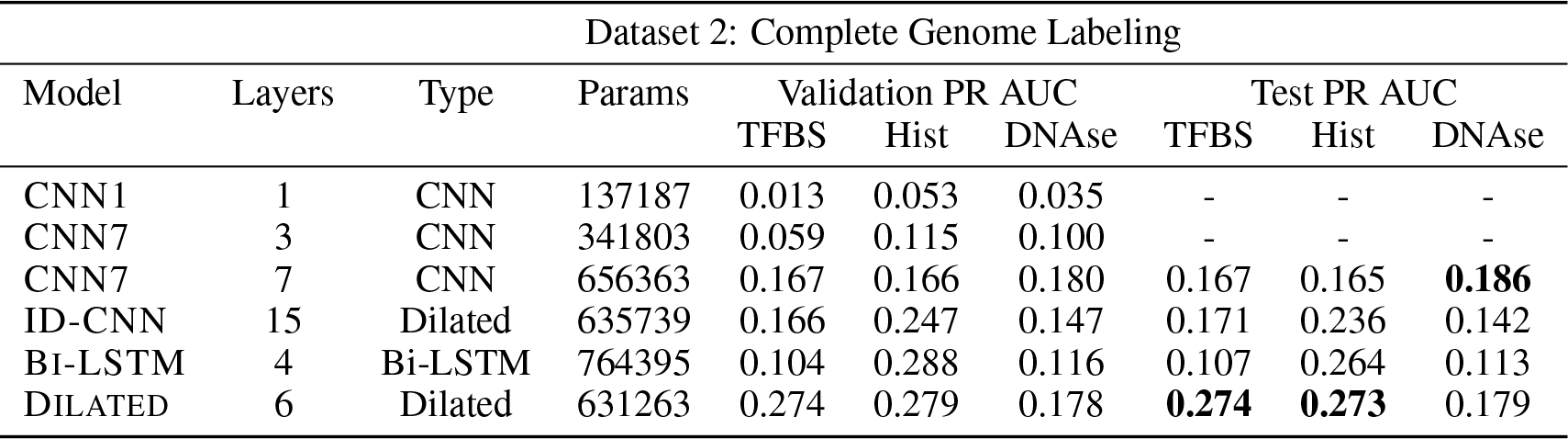
Models for Dataset 2 and Precision-Recall Area Under Curve (PR AUC) scores. We report the number of parameters in the best-performing hyperparameter configuration. ID-CNN is the model from Strubell et al. (2017). We see much higher performance using dilated convolutions on predicting transcription factor binding sites and histone modifications, but no improvement on predicting DNAse hypersensitivity sites. Note that this because this is a new dense prediction task, these results should not be directly compared to those in Table 1.

| Dataset 2: Complete Genome Labeling |  |  |  |  |  |  |  |  |  |
| --- | --- | --- | --- | --- | --- | --- | --- | --- | --- |
| Model | Layers | Type | Params | Validation PR AUC |  |  | Test PR AUC |  |  |
|  |  |  |  | TFBS | Hist | DNase | TFBS | Hist | DNase |
| CNN1 | 1 | CNN | 137187 | 0.013 | 0.053 | 0.035 | - | - | - |
| CNN7 | 3 | CNN | 341803 | 0.059 | 0.115 | 0.100 | - | - | - |
| CNN7 | 7 | CNN | 656363 | 0.167 | 0.166 | 0.180 | 0.167 | 0.165 | **0.186** |
| ID-CNN | 15 | Dilated | 635739 | 0.166 | 0.247 | 0.147 | 0.171 | 0.236 | 0.142 |
| Bi-LSTM | 4 | Bi-LSTM | 764395 | 0.104 | 0.288 | 0.116 | 0.107 | 0.264 | 0.113 |
| DILATED | 6 | Dilated | 631263 | 0.274 | 0.279 | 0.178 | **0.274** | **0.273** | 0.179 |

Dilated convolutional models perform the best on both TFBS and histone modification prediction, and do marginally worse than the best non-dilated models on predicting DNAse hypersensitivity sites. This shows that dilated convolutions can be effective at capturing structure in the genome. In contrast, while BI-LSTM performed better than CNN3, it did worse than the convolutions, particularly on TFBS prediction. This suggests that the LSTM architecture is less effective at this task, either due to vanishing gradients or difficulty in learning gated recurrent connections. This suggests that even though LSTMs are effective on short sequences, when trying to capture properties of long genetic sequences, dilated convolutions are an important architecture to consider.

On DNAse hypersensitivity prediction, standard convolutions do well. This may be because accessible regions locally have highly explanatory motifs and having additional context far away does not improve the accuracy, while the other two marker types are more easily characterized with access to distal motifs. This is also consistent with the high performance of both DILATED6 and Bi-LSTM in predicting DNAse sites in Dataset 1 compared to TFBS and histone modifications.

To further investigate the trained models, we visualize receptive fields by sampling a validation sequence and backpropagating an error of 1 from every positive output for a random regulatory marker. In Figure 2, we plot the output locations (blue) and the norm of the error to the inputs (black), which gives a visual representation of the receptive field. We observe that the standard convolution has a narrow receptive field while the dilated convolution has a wider one, as the gradient is high for a wider input in Figure 2(b). In contrast, the LSTM model in Figure 2(c) has gradient backpropagated widely, but it usually has a low magnitude. It does not appear that the LSTM models are able to learn the long-distance dependencies that the dilated model captures, meaning that though LSTM models were successful on short inputs in Dataset 1, they were unable to scale to larger inputs on Dataset 2.

**Figure 2:**
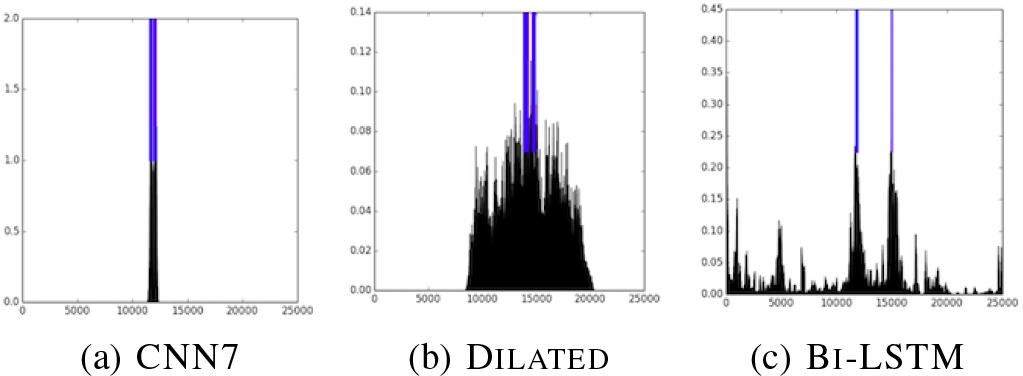
We visualize the norm of the gradient to the inputs (black). This gives an indication about the actual receptive field that was used to make a decision at the outputs (blue). Notably, we see that CNN7 has a narrow receptive field, DILATED has a wide high-magnitude field, and BI-LSTM has a wide but low-magnitude field.

## 4 Conclusion

We introduce a new data set with larger DNA contexts and base pair level data. On this data set, we show that dilated convolutions can outperform both CNNs and LSTMs and appear to capture long-distance relationships in DNA. This suggests that dilated convolutions are an important architecture to consider for genetic modeling. Next, we intend to incorporate more DNA structural information, such as Hi-C data that measures DNA conformation (Belton et al., 2012). We also intend to study whether a hybrid architecture can effectively predict all marker types, particularly the DNAse sites.

## 6 Acknowledgements

We thank David Kelley for his insights on the biological mechanisms behind these regulatory factors, the necessary steps to pre-process ENCODE data, and his general guidance in interpreting the results of this work.

## 7 Supplemental Information: Architecture Details and Hyperparameters

### Dataset 1: Short Sequence Prediction Benchmark

We summarize the high-level architecture that we use for this task in Figure 3 with an example with two convolutional layers. All of the models have the same input representation, and have fully-connected layers at the end. The differences between the models exists between the Input and Flatten layers of Figure 3, as each model uses a different type of convolutional layer and one uses a bidirectional LSTM. Note that in the logistic regression and multilayer perceptron models, we directly flatten the inputs and use fully-connected layers, rather than first applying convolutions.

**Figure 3:**
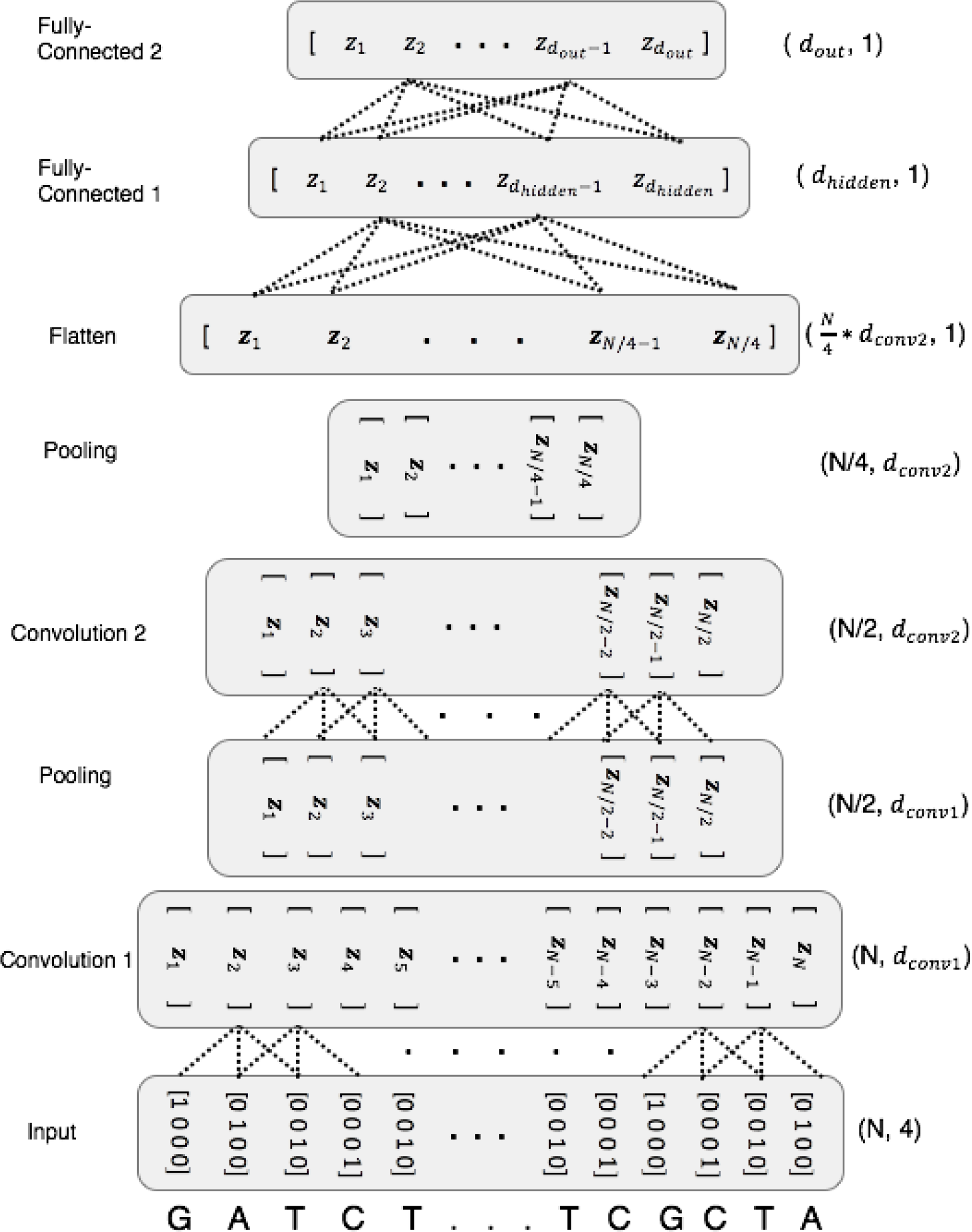
An overview of the architecture we use on Dataset 1. The input (bottom) is a sequence of N nucleotides of DNA, represented as one-hot encoded vectors. The shape of the tensor at each layer is listed on the right. This example shows 2 convolutions, each followed by a pooling layer, and finally two fully-connected layers. Note that there are more convolutions and pooling in some of our models, and we also use dilated convolutions. Additionally, we use a bidirectional LSTM in between the Input and Flatten layers for one of the baseline models. The fully connected layers apply to the flattened input. There are *d*_*out*_ = 919 outputs for each sequence of *N* = 1000 inputs.

We list the hyperparameters we tested in Table 3 in finding the optimum DILATED6 model. We adjusted the number of filters in each layer, represented by {*f*_1_, *f*_2_, *f*_3_}. *f*_1_ and *f*_2_ are for the first two layers and *f*_3_ is for all further layers of the model. We also test the kernel width of the convolution, represented by *k*. Finally, we adjust the size of the final hidden layer, represented by *d*_*hidden*_.

**Table 3:** Hyperparameters tested in Task 1.

| Hyperparameter | Options |
| --- | --- |
| Learning Rate | Between .0001 and .1 |
| Batch Norm Decay Rate | {.9, .99} |
| Dropout | {.1, .01} |
| $k$ | {4, 8, 12} |
| $f_1$ | {120, 240, 360} |
| $f_2$ | {120, 240, 360} |
| $f_3$ | {120, 240, 360} |
| $d_{hidden}$ | {500, 919, 1500} |

### Dataset 2: Complete Genome Labeling with Long-Range Inputs

We summarize the high-level architecture of this task in Figure 4. The differences between the models we implement for this task exists between the Embedding and first Linear (fully-connected) layer of Figure 4. In contrast with Dataset 1, we are now making dense predictions, meaning one set of predictions for every nucleotide in the input sequence, which leads to a different model structure.

**Figure 4:**
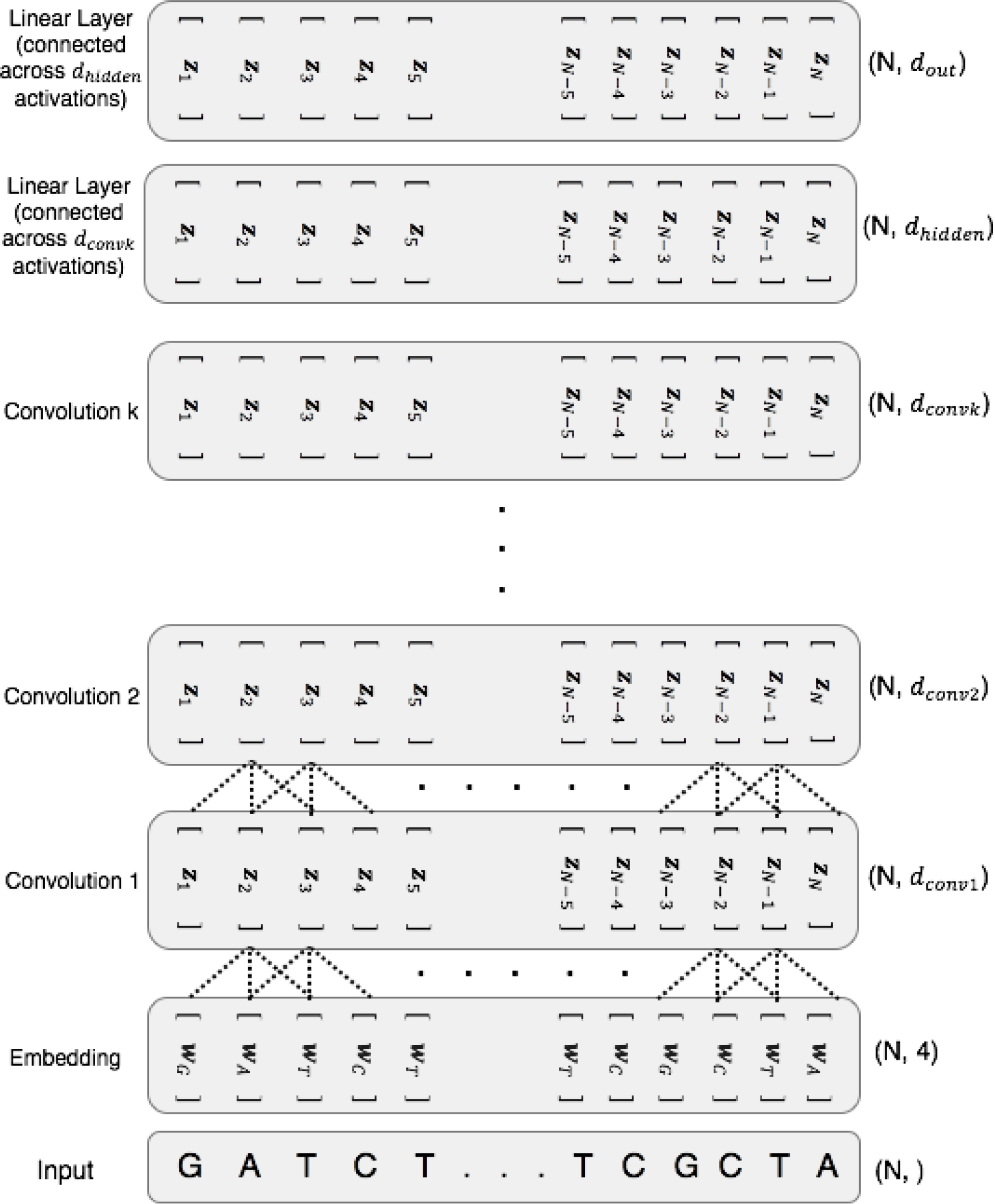
An overview of the convolutional architecture used on Dataset 2. The input (bottom) is a sequence of N nucleotides. The shape of the tensor at each layer is listed on the right. The first layer embeds each element into a higher-dimensional vector space. After that, there are *k* convolutions (or dilated convolutions) with ReLU activations. Furthermore, we test adding pooling layers with a stride of 1 in between convolutional layers. In the LSTM model, the convolutions are replaced with a bidirectional LSTM. Finally, the last two linear layers do not connect elements that are horizontally separated in the above diagram, but only those within the activations for a particular element. So, they apply a linear map to each of the activations 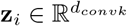. In other words, the first linear layer has weights which are 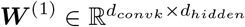, and the second has weights which are 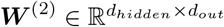. In practice, *N* = 25000, and *d*_*out*_ = 919.

We list the hyperparameters we tested in Table 4. For the convolutions, we adjusted the number of filters in each layer, represented by {*f*_1_, *f*_2_, *f*_3_}. *f*_1_ and *f*_2_ are for the first two layers and *f*_3_ is for all further layers of the model, if applicable. For the LSTM, *f*_1_ is the number of filters in the first convolution, *f*_2_ referred to the state size of each LSTM, and *f*_3_ is the number of filters in the deconvolution. For the LSTM, we also adjust the stride for the first convolution and final deconvolution. Each model had one hidden layer after the convolutions, with *d*_*hidden*_ units.

**Table 4:** Hyperparameters tested in Task 2.

| Hyperparameter | Conv Options | LSTM Options |
| --- | --- | --- |
| Learning Rate | [.001, .1] | [.001 and .1] |
| Batch Decay | {.9, .99} | {.9, .99} |
| Dropout | {.1, .01} | {.1, .01} |
| $f_1$ | { 64, 128 } | 128 |
| $f_2$ | {120, 240} | {20, 40, 80} |
| $f_3$ | {50, 128} | {50, 128} |
| Stride | 10 | {10, 20, 50, 100} |
| $d_{hidden}$ | {125, 200} | {64, 125, 200} |

